# Patchy Perception: Rethinking Eye Movements through the Lens of Foraging Theory

**DOI:** 10.1101/2025.03.20.644341

**Authors:** Tal Nahari, Ahmed El Hady

**Affiliations:** Affective Brain Lab, Department of Experimental Psychology, University College London, London, UK; Max Planck UCL Centre for Computational Psychiatry and Ageing Research, University College London, London, UK; Department of Collective Behavior, Max Planck Institute of Animal Behavior, Konstanz, Germany; Department of Biology, University of Konstanz, Konstanz, Germany; Centre for Advanced Study of Collective Behavior, University of Konstanz, Konstanz, Germany

## Abstract

Foraging theory provides a powerful framework for reframing visual attention mechanisms, conceptualizing eye movements and visual search as specialized instances of patch foraging problems, rather than viewing them solely through traditional cognitive psychology paradigms. This approach offers new insights into how visual attention optimizes exploration strategies in humans and animals. Using human data from image exploration tasks with items held in short-term memory, we demonstrate that participants spend more time on informationally rich stimuli (memory-matched images) and employ strategies to minimize travel time to these high-value targets—behaviors consistent with optimal patch foraging. Our proposed analytical approach aims at a more foundational question in movement ecology: Why might animals partition their environment into patches of information as they move through it? This work provides a foundation for new ways to analyze already existing data and design experimental paradigms bridging visual neuroscience and behavioral ecology approaches.

## INTRODUCTION

Foraging is a foundational decision-making process that all animals perform in order to survive in their environment. Behavioral ecology is rich with quantitative studies of foraging behaviors, especially “rules of thumb” that animals use to forage efficiently in their environments. One of the most well studied foraging behaviors is patch foraging. In its simplest form, patch foraging is when an animal enters a patch of food, harvests resources, and then leaves to search for another patch of food ([1]). While natural environments rarely exhibit the idealized patchiness of theoretical models, the conceptualization of resources as spatially clustered provides a valuable formalism for differentiating between local exploitation and global exploration phases. This framework enables quantitative characterization of multiple behavioral parameters, including patch residence time, resource consumption rates, and inter-patch movement trajectories. An open question in sensory ecology is whether animals perceive the environments they are living in as patches of differential information content? Would such a sensory strategy optimize animal movement and foraging in uncertain environments? A way to approach these questions might arrive from cognitive science. Cognitive systems are not merely tuned to tangible foraging for food resources in the environment. Rather, evolutionary pressures may have shaped cognitive processes to reflect information-seeking patterns analogous to those observed in ecological contexts, thus providing a general purpose cognitive architecture that can be used both for information seeking or food gathering ([2]). This alignment suggests that the principles governing ecological movement in physical spaces may also apply to the abstract landscapes of cognition, and will be evident in other domains of search, either mentally (i.e, searching for creative solutions; [3]) or sensory, i.e, via shifts of gaze.

Examining search behaviors in the visual system is particularly interesting, since unlike other sensory systems, vision operates predominantly through active sampling, where the eyes continually shift gaze to acquire information—a process known as the active vision loop ([4]). This dynamic interplay between what we look at and what interests us creates a cascade of mutual influence ([5]), shaping both perception and behavior. We look at what conveys more information; and we also attribute more information and value to what we look at ([6]). But what characterizes visual exploration behavior? Since the early days of eye-tracking research, distinct movement patterns have been observed, separating between the time where the eyes are fixated on a single location (coined fixations) and the rapid shifts in gaze, known as saccades ([7]). Intriguingly, fixations tend to stabilize on locations rich in information—whether that information is low-level visual features, such as density or brightness ([8]), or semantically meaningful content, such as attractiveness ([9]), familiarity ([10–12]) or short-term memory ([13]). This behavior highlights the inherent efficiency of visual search in directing attention toward valuable informational patches (see figure 1). These visual search dynamics are reminiscent of the way animals forage in their environments. The eyes, similarly to animals, are drawn to areas with higher information or resource value. Could it be that one mechanism drives both the eye movement patterns and the animal foraging behaviors? If so, what could be such mechanisms? Behavioral ecology has proposed various rules to explain animal foraging behavior, with the most influential being the Marginal Value Theorem (MVT). MVT states that an animal can optimize energy intake by leaving its current patch when the current reward rate falls below the global average reward rate of the environment ([14]). This gives a theoretical foundation for assessing optimal decision-making, which has also been validated in many behavioral studies ([15–20]). Recently, a series of mechanistic models have been proposed in order to account for the potential cognitive mechanisms animals might be employing to implement MVT or MVT-like rule ([21]). The mechanism proposed by these models is that the animal accumulates evidence as it enters the patch till it reaches a threshold then leaves. Similar models are used in the field of experimental psychology, the drift diffusion model formalism used in cognitive neuroscience of decision-making behaviors ([22]). Interestingly, Drift diffusion models have been used in eye tracking experiments for value-based decisions before ([23]). Here, we provide preliminary evidence for integrating canonical movement models—traditionally grounded in physical and physiological principles of animal behavior—with human oculomotor data. We analyze a dataset ([24]) to examine gaze transitions across spatial locations and fixation durations within interest areas, comparing patterns associated with faces stored in short-term memory (targets) versus non-target stimuli. This analysis establishes preliminary evidence for conceptual parallels between patch foraging theory and human visual exploration, suggesting a unified governing mechanism underlying fundamental search processes.

**FIG. 1.**
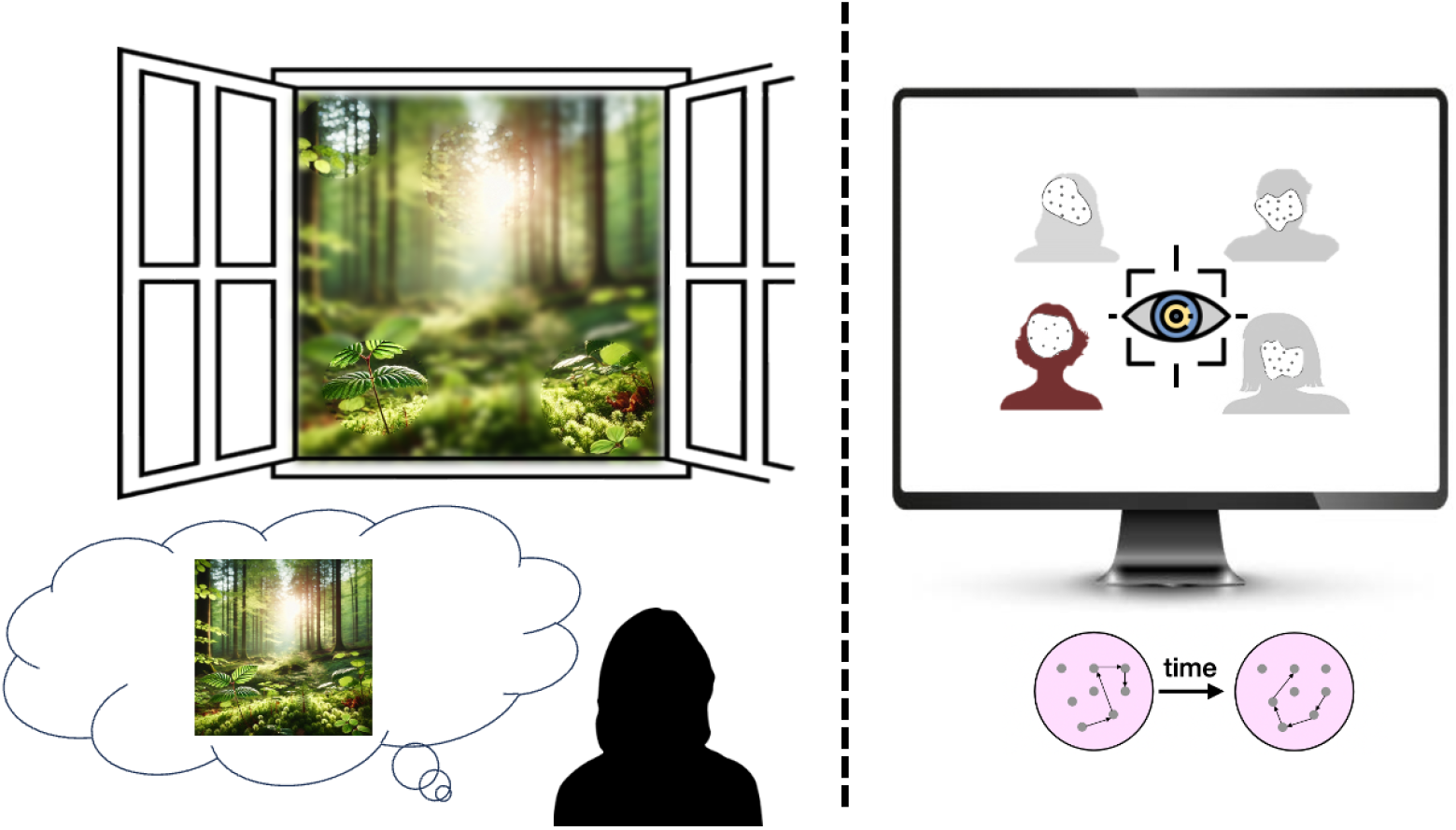
Visual exploration and eye movement analysis. (A) Selective visual attention during natural scene viewing. The foveal region provides high-resolution information while peripheral vision remains low-resolution, despite subjective perception of scene completeness. (B) Example paradigms for quantifying visual exploration behavior. Eye-tracking methodology enables systematic measurement of gaze patterns in response to manipulated visual stimuli, conceptualizing visual search as a spatial foraging problem.

## METHODS

### Experimental Methods

All experiments were approved by the ethics committee of the social science faculty in the Hebrew University of Jerusalem (Jerusalem, Israel).

### Participants

The sample included 59 university students (32 women; Average age 27.7, sd 3.9). After exclusion (see criteria below), the sample consisted of 48 participants with normal or corrected-normal vision and valid eye movements data from the entire experiment. All participants signed an informed consent before the experiment. They were granted either course credits or 40 NIS (approminately 10 dollars). The experiment was approved by the psychology department ethics committee.

### Stimuli

We used 64 pictures of past years’ students of the Hebrew University that were held in the University database. All pictures were of neutral expressions, front facing the camera. We normalized the pictures for brightness using a Matlab code (find the code in: https://osf.io/wgfb4/). The stimuli were displayed on a 24 inches BenQ 3d monitor, with a 120-Hz refresh rate BenQ monitor and a 1024 × 768 screen resolution, corresponding to a screen size of 47.6 × 28, situated at a distance of 60 cm from the participants’ eyes.

### Memory task

The short-term memory task included two blocks in random order, one with 64 trials of the single-first condition and the other with 64 trials of the multiple-first condition. Here, we focus only on the single-first condition, as we want to see how search alters in multiple interest areas in which one image contains more information (see Figure 2). Each trial began with a drift check, allowing a deviation of only 0.75 degree of visual angle between the predicted gaze position and the center of fixation point. Larger deviations were accompanied by an error beep and led to a repeated calibration process. Participants saw a single face (2000 ms), followed by a delay with a blank screen and a fixation cross (3000 ms), a multiple display of four faces (5000 ms) and a blank screen with a central fixation point (1000 ms). During the multiple display, participants were required to press one of two keys indicating whether one of the faces had been presented in the previous single display or not. Parallel displays were displayed longer as they consisted of more faces, leaving more observation time for each face. The faces in both displays (i.e., the multiple and the single ones) were from the same sex, and the correct answer in half of the trials was “yes” and in the other half “no”, in a random order in both conditions. At the end of the experiment, participants were rewarded based on their accuracy – if they were correct (accurately reporting if a face in the retrieval display was also displayed in the encoding display) in over 90 percent of the trials, they were awarded a bonus (10 NIS, approximately 2.5 dollars). Participants performed a practice session before the block, and had to complete at least three correct practice trials out of five, otherwise, they underwent another session of five training trials. Based on the final debriefing, and similarly to our previous studies ([11, 25]), If more than 16 pictures were removed from the data of a single participant (equivalent to a quarter of the total number of pictures), or more than 1 standard deviation above the mean of the rest of the participants, the data of the participant was excluded from the analysis (11 out of 59 participants). The final sample consisted of 48 participants.

**FIG. 2.**
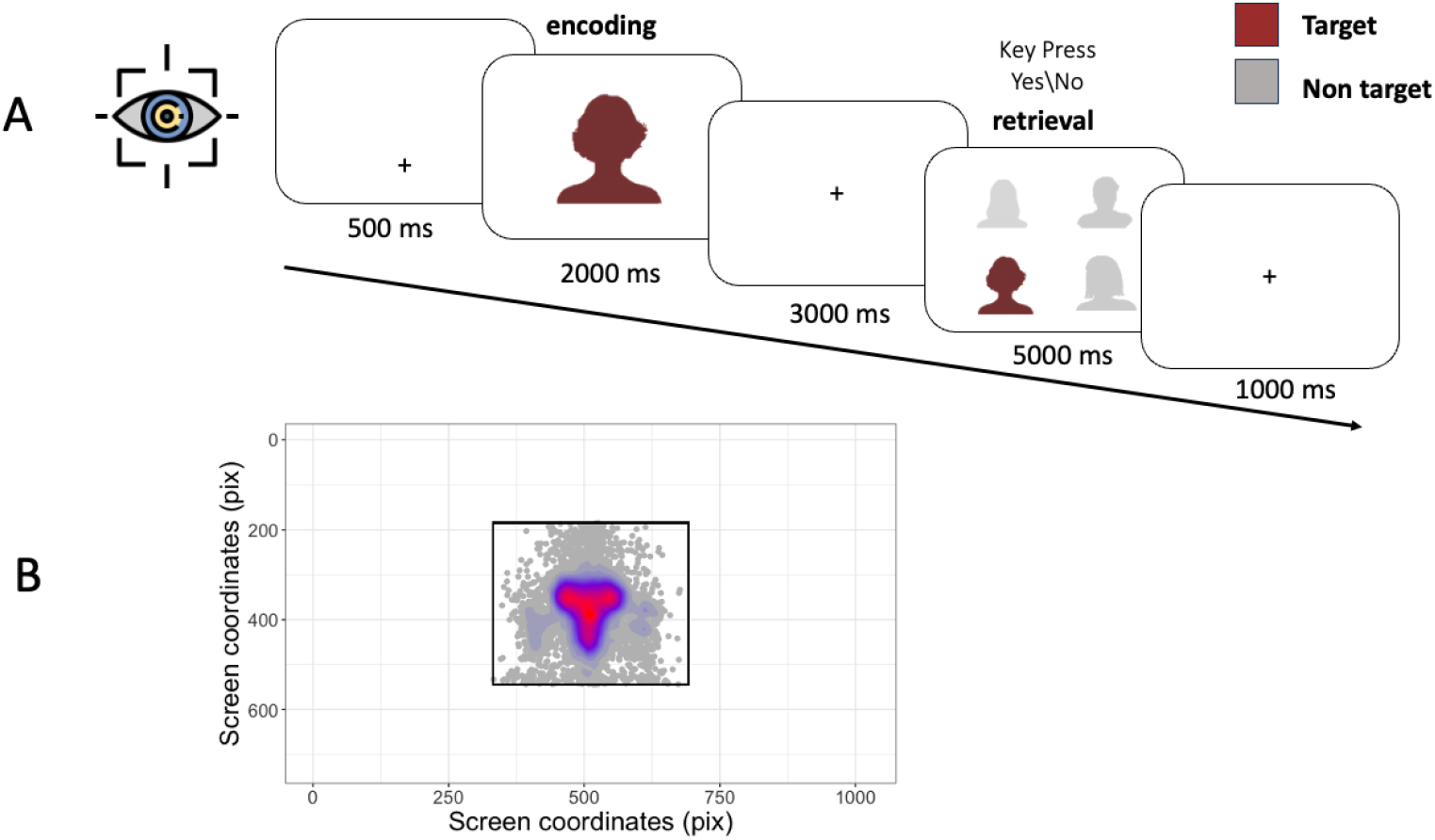
Experimental design and visual attential analysis. (A) Sequential recognition paradigm. Partici-pants encoded a single target face during the initial presentation phase, followed by a two-alternative forced-choice recognition task (to indicate whether the target is present - yes or no). (B) Spatial distribution of visual fixations during the encoding phase. The superimposed heat map illustrates fixation density, with rectangular boundaries demarcating the stimulus presentation area.

### Eye Tracking

The experiment began with a standard 9-point calibration and validation procedure provided by Eyelink 1000+ (SR Research Ltd., Mississauga, Ontario, Canada). Average accuracy in the validation procedure ranged between 0.25º - 0.82º of visual angle. In each trial, four identical rectangle interest areas (size: 360 on 360 pixels; 16.7 on 16.7) surrounded each one of the four faces in the multiple display, separated horizontally by 55 pixels (2.5) and vertically by 48 pixels (2.23). In the single display, an interest area was outlined around the presented face (size: 480 on 480 pixels; 22.3 on 22.3).

### Analysis Methods

#### Fixation parsing

The eye-tracking measures are based on EyeLink’s standard parser configuration: samples were defined as a saccade when the deviation of consecutive samples exceeded 30 °/s velocity or 8,000 °/s^2^ acceleration. Samples gathered from time intervals between saccades were defined as fixations.

#### Spatial analysis

For the spatial analysis, we calculated an average within each subject within each of the interest areas.

#### Duration analysis

The time to reach the location was calculated by the first fixations on the location, depending on whether that location contained a target item or not. All fixations were averaged within subject, trial, and location, and then averaged across locations to test for statistical significance (see Figure 3). Similarly, we analysed the mean fixation duration on each location by averaging within subject, trial and location, and then across locations.

**FIG. 3.**
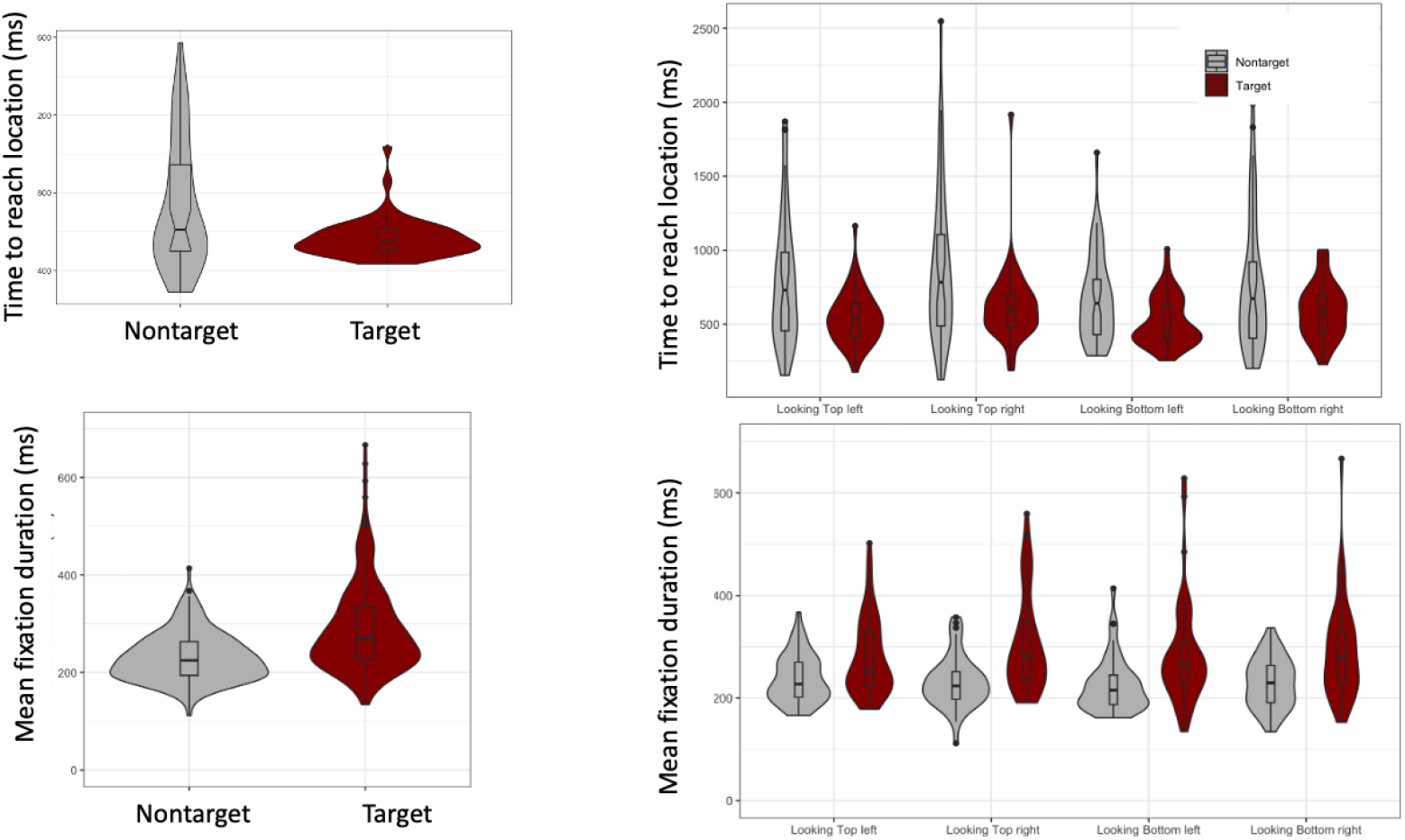
Temporal dynamics of visual search behavior as a function of stimulus type. (A) time to reach the target location across spatial positions. Participants demonstrated significantly reduced latency when navigating to positions containing target stimuli compared to non-target positions, consistent across all spatial locations. (B) Fixation duration analysis. Mean fixation durations were significantly prolonged for target stimuli relative to non-target stimuli, indicating enhanced attentional engagement with task-relevant information. Violin plots illustrate the complete distribution of values across all subjects, with width indicating data density at each point. Embedded boxplots show the interquartile range (25th-75th percentiles) with the horizontal line denoting the median value.

## RESULTS

Since the accuracy rates are at ceiling (t(48) = 89.28, p*<*0.001, d = 12.75), we used all the trials in the task. During encoding, participants scanned the single face in a manner that replicated previous findings in the literature on face perception ([26]) - spending most of the time on the eyes and mouth (see figure 2B for a heat map of all participants). We then turned to analyze the multiple face display as a patch foraging problem. The four images, each with its own interest area, are equivalent to four patches. we hypothesized the following:

1. The subjects would spend more time on the short-term memory stored image akin to patch residence time where foragers will spend more time in highly informative patches compared to the ones with lower information rates.
2. The subjects will adopt a search strategy that minimizes the arrival time to the most informative image. Here, the target item, stored in short-term memory.

Our analysis focused first on the time it took to reach each area of interest. Participants demonstrated significantly reduced latency when navigating to positions containing the target face previously encoded in short-term memory, regardless of spatial position (t(47)=3.27, p*<*0.001). This latency is akin to the arrival time to the highest rewarding patches. Furthermore, mean fixation durations were significantly prolonged for target stimuli relative to non-target stimuli (t(47) = 5.43, p *<* 0.001), suggesting enhanced attentional engagement with memory-matching stimuli (Figure 3). Mean fixation durations here are a proxy for patch residence times.

To examine spatial visual search patterns in addition to temporal dynamics, we analyzed participants’ gaze allocation across the display (Figure 4). We then quantified the fixational transitions preceding response execution, focusing exclusively on target-present trials. The analysis incorporated the four spatial quadrants (top-right, top-left, bottom-right, and bottom-left) to characterize the transitional flow between regions during the search process. This is akin to animals moving and searching between patches searching for the most rewarding one (Figure 5).

**FIG. 4.**
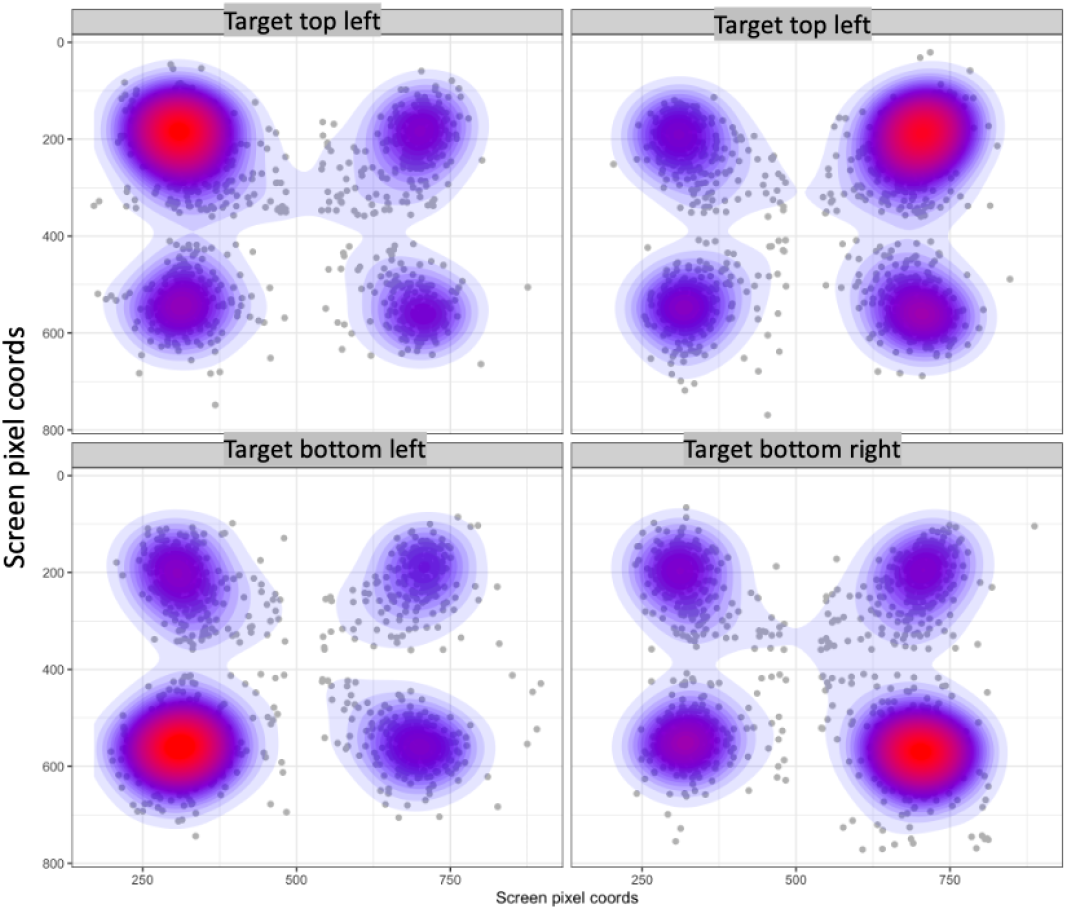
Spatial distribution of visual fixations during the retrieval phase as a function of target location. The superimposed heat map illustrates fixation density.

**FIG. 5.**
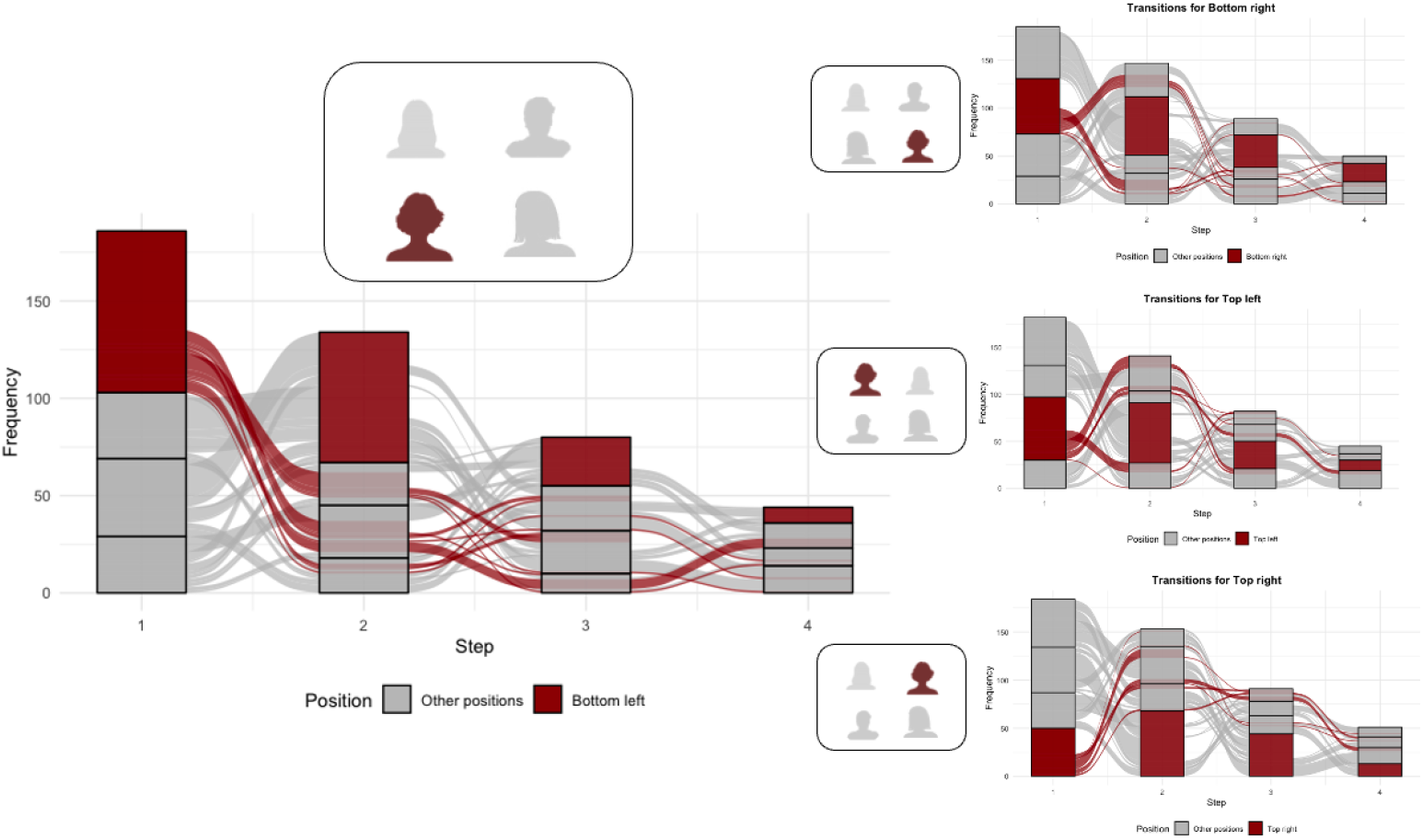
Sequential eye movement patterns as a function of target location. The visualization depicts the temporal flow of visual attention across defined spatial regions during the search task before the key press, depending on different locations of the target. Bar height indicates fixation frequency at each location prior to response execution. The alluvial streams illustrate the sequential progression of gaze patterns, demonstrating how visual exploration strategies are modulated by target position

To evaluate whether search termination corresponded with target location, we employed a Wilcoxon signed-rank test with continuity correction. Results revealed a statistically significant relationship (V= 31, p = 0.039), indicating that participants’ final fixations were disproportionately directed toward target locations compared to what would be expected by chance allocation. This finding suggests that memory-guided search effectively biases visual exploration toward behaviorally relevant stimuli. They end up preferentially at the most informative familiar images through a variety of visual search sequences. The general search pattern is spending more time at the more informative image, despite the variable search trajectories (Figure 4). Taken together, participants reach the target more quickly, and spend longer fixation duration on it. Moreover, they transition towards the target location in less steps, and finish their searches on the target location.

## DISCUSSION

In this study, we propose an innovative framework for understanding visual exploration through the lens of patch foraging theory, particularly when examining items stored in short-term memory. While previous research has directly applied foraging paradigms to eye movement studies ([27, 28]), we argue that many other tasks involving recorded eye movements can be conceptualized within a patch foraging framework, thereby expanding the analytical scope of visual cognition research and improving the theoretical search models. Our findings support our central hypothesis: participants demonstrated significantly reduced latency when navigating to target faces stored in short-term memory, regardless of spatial position, and exhibited prolonged fixation durations on these targets. However, substantial individual differences were observed in exploration patterns. Notably, participants did not universally fixate on the target first before response execution, and in numerous instances, continued to examine alternative stimuli even after initial target identification. This variability suggests that while memory representations guide visual search, the process is not deterministic but rather probabilistic in nature. The general underlying mechanism might be minimizing the arrival time to the most informative image, a foraging strategy which has been shown to be bayes optimal in the case of two or more non - depleting resource patches ([29]). Despite the promising results of our study, several limitations warrant consideration. First, our experimental paradigm employed static visual arrays with predefined target categories, which may not fully capture the dynamic and continuous nature of real-world visual foraging. Second, while we observed consistent patterns of memory-guided search behavior, our trial sample size limited comprehensive exploration of individual differences that might reveal distinct foraging strategies. Additionally, the laboratory setting inevitably introduces artificial constraints that may not generalize to naturalistic contexts where visual search occurs amid competing goals and distractors of varying ecological relevance. Finally, although our framework provides a conceptual bridge between visual search and patch foraging, further computational modeling is needed to formalize the precise decision mechanisms governing the observed exploration patterns. Still, these findings have implications that extend beyond short-term memory paradigms, and are worthy of dedicated task designs. For instance, researchers could implement parametric memory manipulations or pairing with reward associations to achieve more nuanced modulations of stored information representations, to see the dynamic between different quantities of information. In addition, environmental characteristics could be systematically varied to introduce different levels of perceptual certainty or ambiguity. This will allow fine-grained analysis that will verify specific predictions, for example - whether subjects move across images weighing their informational content and adjusting their residence time in it like in the case of patch foraging. Furthermore, this framework could be productively extended to investigations of search within semantic networks. Research has demonstrated that human cognitive processes exhibit auto-correlated structures wherein information clusters into “patches,” analogous to resource distributions in natural environments ([2, 3]). Such paradigms would likely elicit comparable foraging behaviors, enhancing our understanding of how cognitive systems navigate both physical space and abstract informational landscapes. Moreover, given the ubiquity of measuring eye movement in neuroscientific and psychological behavioral paradigms, a reanalysis of these data within our proposed framework can contribute significantly to our understanding of how animals forage within the information space of their perceived environments. In conclusion, extending patch foraging models across species and domains—encompassing both physical and abstract information search—offers considerable potential for elucidating the overarching exploration mechanisms that govern cognitive systems and facilitate resource acquisition in diverse environments. By integrating theoretical perspectives from movement ecology, visual cognition, and computational modeling, we can develop a more comprehensive framework for understanding behavioral manifestations of fundamental cognitive processes and their underlying neural mechanisms.

## AVAILABILITY OF DATA AND MATERIALS

The datasets generated and/or analyzed during the current study are not publicly available, but are available from the corresponding author on reasonable request.

## COMPETING INTERESTS

The authors declare that they have no competing interests.

## ACKNOWLEDGMENTS

The idea of this project was born at NeuroBridges 2023 school in Cluny, France. We would like to thank Prof. Yoni Pertzov from the Hebrew University of Jerusalem for allowing us to reuse the data and materials. TN would like to thank Prof. Luca Giuggioli for his support in the initial ideation. AEH is supported by the Deutsche Forschungsgemeinschaft (DFG, German Research Foundation) under Germany’s Excellence Strategy EXC 2117422037984.

## Notes

### Competing Interest Statement

The authors have declared no competing interest.

